# The cortical bone metabolome of C57BL/6J mice is sexually dimorphic

**DOI:** 10.1101/2021.08.06.455423

**Authors:** Hope D. Welhaven, Ghazal Vahidi, Seth T. Walk, Brian Bothner, Stephen A. Martin, Chelsea M. Heveran, Ronald K. June

**Affiliations:** Department of Chemistry & Biochemistry, Montana State University, Bozeman, MT 59717; Molecular Biosciences Program, Montana State University, Bozeman, MT 59717; Department of Mechanical & Industrial Engineering, Montana State University, Bozeman, MT, 59717; Department of Microbiology and Cell Biology, Montana State University, Bozeman, MT, 59717; Translational Biomarkers Core Laboratory, Montana State University, Bozeman, MT, 59717

**Author notes:** Co-corresponding Authors, Chelsea M. Heveran, Ph.D., Dept. of Mechanical & Industrial Engineering, Montana State University, PO Box 173800, Bozeman, MT 59717-3800, Ronald K. June, Ph.D., Dept. of Mechanical & Industrial Engineering, Montana State University, PO Box 173800, Bozeman, MT 59717-3800.

**Keywords:** metabolomics, metabolism, sex differences, bone quality

## Abstract

Cortical bone quality, which is sexually dimorphic, depends on bone turnover and therefore the activities of remodeling bone cells. However, sex differences in cortical bone metabolism are not yet defined. Adding to the uncertainty about cortical bone metabolism, the metabolomes of whole bone, isolated cortical bone without marrow, and bone marrow have not been compared. We hypothesized that the metabolome of isolated cortical bone would be distinct from that of bone marrow and would reveal sex differences. Metabolite profiles from LC-MS of whole bone, isolated cortical bone, and bone marrow were generated from humeri from 20-week-old female C57Bl/6J mice. The cortical bone metabolomes were then compared for 20-week-old female and male C57Bl/6J mice. Femurs from male and female mice were evaluated for flexural material properties and were then categorized into bone strength groups. The metabolome of isolated cortical bone was distinct from both whole bone and bone marrow. We also found sex differences in the isolated cortical bone metabolome. Based on metabolite pathway analysis, females had higher lipid metabolism, and males had higher amino acid metabolism. High-strength bones, regardless of sex, had greater tryptophan and purine metabolism. For males, high strength bones had upregulated nucleotide metabolism, whereas lower strength bones had greater pentose phosphate pathway metabolism. Since the higher strength groups (females compared with males, high strength males compared with lower strength males) had higher serum CTX1/P1NP, we estimate that the metabolomic signature of bone strength in our study at least partially reflects differences in bone turnover. These data provide novel insight into bone bioenergetics and the sexual dimorphic nature of bone material properties in C57Bl/6 mice.

## 1. Introduction

Sex differences exist in mouse bone tissue across several length scales ^(1–16)^. In C57Bl/6 mice, females and males frequently exhibit different whole bone material properties ^(1,5,6,8,10)^ Females can have statistically significantly higher bone strength ^(5)^, although more commonly studies report data that suggest that females have increased strength for females but do not specifically test sex differences ^(1,5,10)^. In other work with BALB/c mice, 6 months old females have higher modulus and tougher femurs than males, but femur strength does not differ ^(17)^. Sex differences are seen in Wistar rats as well, with females having higher mineral:matrix from Raman spectroscopy ^(18)^. Sex differences in bone strength also exist for humans, although unlike mice, young adult males have stronger bones than young adult females ^(19,20)^. There are also sex differences in remodeling bone cell populations. For instance, female rodents often have a higher osteoclast surface ^(11–15)^ and a higher osteoblast surface ^(12–16)^. Bone formation and resorption together influence bone quality ^(21,22)^. Because bone formation and resorption both require cellular energy production and utilization ^(23,24)^, the metabolic products of bone cells are of high value in improving our understanding of the basis of sex differences in cortical bone quality.

Metabolomics, the study of small molecule intermediates called metabolites, has the potential to provide novel insight into the connection between bone cell metabolism and bone quality. Bone cell metabolism is currently studied using several different approaches. The Seahorse assay is commonly applied to bone marrow to determine oxygen consumption rate ^(3,25–30)^. However, cortical bone and its metabolism likely differs from bone marrow. Shum *et al* used an LC-MS approach to study marrow-flushed cortical mouse bone. They found a glycolytic shift from 3-months to 13*months of age in male C57Bl/6 mice. These results helped to contextualize changes to mitochondrial function with age ^(31)^. In another study, Zhao *et al* used LC-MS marrow flushed mouse femurs to understand changes with OVX to cortical bone bioenergetics. They found that OVX mice had disordered lipid and amino acid metabolism ^(32)^. Collectively, these prior studies show that cortical bone metabolism changes in aging and disease models. However, it is still unknown how sex influences the cortical bone metabolome.

Another important question is whether, and how, cortical bone metabolism differs from bone marrow. Bone marrow contains mesenchymal stem cells which give rise to numerous cell types including adipocytes, osteoblasts, osteocytes, as well as myeloid and immune cells^(33)^. Cortical bone is mostly cellularized by osteocytes (~90%) ^(34,35)^ but metabolites may persist in this tissue from bone marrow. This question can be addressed by comparing the metabolomes of isolated cortical bone, bone marrow, and whole bone using metabolomics.

The purpose of this study was to (1) evaluate differences in the metabolomes of whole bone (i.e., cortical and trabecular bone and marrow), isolated cortical bone, and bone marrow (2) assess if the cortical bone metabolome is sexually dimorphic, and (3) determine if metabolomic differences correspond with whole bone strength, a sexually dimorphic cortical bone material property.

## 2. Methods

### 2.1 Animals

Two sets of mice were utilized for this study (total n = 30). First, C57Bl/6J female mice (n=10) were utilized to compare the metabolomes of whole bone, isolated cortical bone, and bone marrow. These mice were purchased from Jackson Labs and acclimated to the Montana State University animal facility for 3 weeks. Second, twenty C57Bl/6J mice (female, n = 10; male, n = 10) were utilized to evaluate sex differences in the cortical bone metabolome. These mice were born and raised at Montana State University.

All mice were housed in cages of 3-5 mice and fed a standard chow fed diet *ad libitum* (PicoLab Rodent Diet 20, 20% protein). All 30 mice were euthanized via cervical dislocation at age 20-21 weeks. All animal procedures were approved by the Institutional Animal Care and Use Committee at Montana State University. Investigators remained blinded to mouse sex during data collection and analyses.

### 2.2 Metabolomics

#### 2.2.1 Experimental design to assess metabolic differences in bone and marrow

We first evaluated differences in metabolomic profiles of isolated cortical bone, bone marrow, and whole bone for humeri from female mice (n = 10). To isolate bone marrow from cortical bone, the distal and proximal ends of right humeri were removed and the cortical shaft flushed with phosphate-buffered saline (PBS). Both cortical bone and marrow were preserved for analyses. The left humerus remained whole.

To extract bone metabolites, whole humeri and isolated cortical bone were placed in liquid nitrogen for 2 hours and pulverized to a powder using an autoclaved aluminum cold sink, rods, and a hammer to optimize extraction. Bone powder was extracted with 3:1 methanol:acetone. Samples were subjected to 5 cycles, each consisting of 1 minute of vortexing followed by macromolecule precipitation at −20°C for 4 minutes. Samples were stored overnight at −20°C to allow for precipitation of remaining macromolecules. The next day, samples were centrifuged to remove cells and debris, supernatant containing metabolites was collected, and dried down via vacuum concentration. Once the supernatant was isolated and dried, metabolites were resuspended with 1:1 acetonitrile:water (Figure 1).

**Figure 1.**
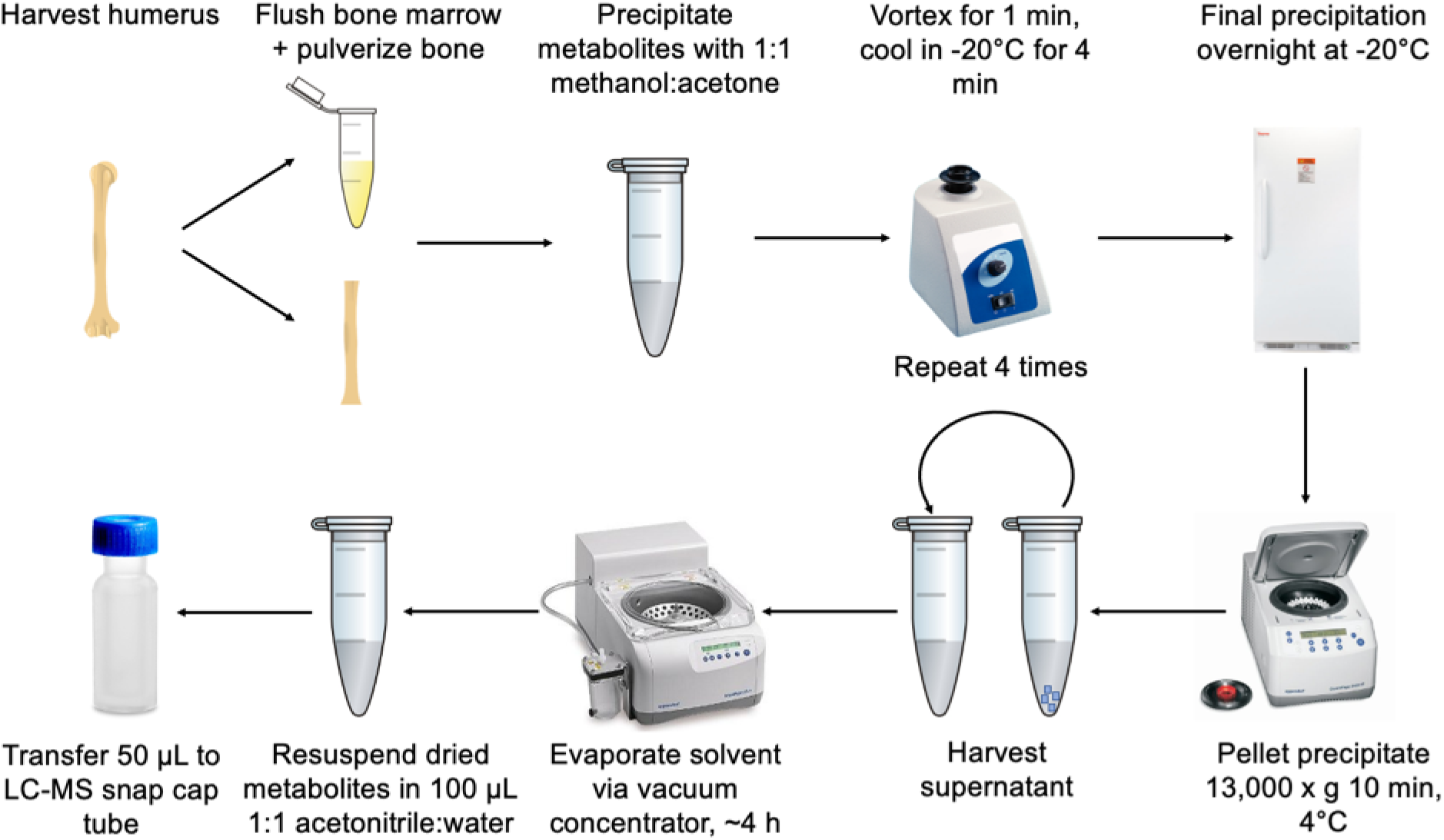
Experimental protocol for extracting metabolites from bone and bone marrow.

Flushed bone marrow underwent a similar extraction protocol. To extract bone marrow metabolites, 1 mL of 70:30 methanol:acetone was added and the mixture was subjected to 5 cycles of vortexing and −20°C precipitation. Samples were then stored overnight at −20°C to precipitate any remaining macromolecules. The next day, samples were centrifuged and supernatant containing metabolites was dried down via vacuum concentration. Dried metabolites were resuspended with 1:1 acetonitrile:water (Figure 1). All solvents used were HPLC-grade or higher.

#### 2.2.2 Evaluation of sex differences in isolated cortical humerus metabolomic profiles

We investigated sex differences in metabolomic profiles of cortical humeri for C57Bl/6 mice (females, n = 10; males, n = 10). Left humeri were dissected, cleaned, and bone marrow flushed to isolate cortical bone tissue. Bones were stored in PBS-dampened gauze at −20°C after dissection until the metabolite extraction. For isolated bone metabolite extraction used the protocol described in 2.2.1.

Following metabolite extraction of whole bone, bone marrow, or isolated bone, all samples (n = 30) were analyzed by Liquid Chromatography Mass Spectrometry (LCMS) using an Agilent 1290 LC coupled to an Agilent 6538 Quadrupole-Time of Flight (Q-TOF) mass spectrometer in positive mode (resolution: ~25,000 FWHM, accuracy: ± 5 ppm). A Cogent Diamond Hydride HIILIC chromatography column was utilized (2.2 μM, 120 Å, 150 mm x 2.1 mm, MicroSolv Leland, NC, United States). 5 μL of sample was injected and blank samples were analyzed every 7-10 samples for quality control to prevent spectral drift and contamination. Agilent Masshunter Qualitative Analysis software was used to identify and export peak intensity values for m/z values in the experimental sample set. LCMS data was then exported and converted for analysis using XCMS. Differential analysis of mass features was completed using MetaboAnalyst. Kyoto Encyclopedia of Genes and Genomes (KEGG) was utilized to confirm retention time and exact mass of detected metabolite features.

### 2.3 Bone marrow adiposity

Proximal tibiae were dissected and fixed for 18 hours in 10% neutral buffered formalin. Tibiae were then decalcified with EDTA, dehydrated in a graded ethanol series, and embedded in paraffin. 5 μm sections were cut longitudinally onto glass slides and stained with hematoxylin and eosin (H&E) per standard protocols.

Sections were imaged using a Nikon Eclipse E-800 and Universal Imaging Corporation’s MetaVue software (version 7.4.6). Images were taken with a 4x objective (909 pixels / 1 mm). A central section proximal tibia was selected for analyses. Adipocytes were manually segmented, counted, and measured using the iPad app YouDoodle and custom MATLAB code. Measurements included mean adipocyte area (mm^2^), marrow cavity area (mm^2^), and adipocyte number density (number of adipocytes per marrow cavity area).

To confirm that the technique utilized to flush marrow from the humerus, additional humeri were assessed. These bones were dissected, harvested, and sections were prepared, imaged, and analyzed consistent with the description for tibiae. Microscopy confirmed that marrow was almost entirely removed from the flushing step (Supplementary Figure 1).

### 2.4 Serum chemistry analysis

Mouse serum was collected at the time of euthanasia via cardiac puncture. Serum was aliquoted and stored at −80°C. Serum was tested for two biomarkers, P1NP and CTX1, using commercially-available kits (MyBioSource, P1NP = MBS703389, CTX1 = MBS722404) via ELISAs using a BioTek spectrophotometer.

### 2.5 Trabecular microarchitecture and cortical geometry

After dissection, left femora were stored at −20°C wrapped in PBS-soaked gauze. These were thawed for MicroCT (Scanco uCT40) analysis and afterwards refrozen as before until flexural testing. Scans were acquired using a 10 μm^3^ isotropic voxel size, 70 kVP, 114 μA, 200 ms integration time, and were subjected to Gaussian filtration and segmentation. Image acquisition and analysis adhered to the JBMR guidelines ^(36)^. Trabecular bone microarchitecture of the distal femur metaphysis was evaluated in a region beginning 200 μm superior to the top of the distal growth plate and extending proximally 1500 μm proximally. The trabecular region was identified by manually contouring the endocortical region of the bone. Trabecular bone was segmented from soft tissue using a threshold of 375 mgHA/cm^3^. Measurements included trabecular bone volume fraction (Tb.BV/TV, %), trabecular bone mineral density (Tb. BMD, mgHA/cm^3^), connectivity density (Conn.D, 1/mm^3^), structural model index (SMI), specific bone surface (BS/BV, mm^2^/mm^3^), trabecular thickness (Tb.Th, mm), trabecular number (Tb.N, mm-^1^), and trabecular separation (Tb.Sp, mm). Cortical bone geometry was assessed in 50 transverse μCT slices (500 μm long region) at the femoral mid-diaphysis and the region included the entire outer most edge of the cortex. Cortical bone was segmented using a fixed threshold of 700 mgHA/cm^3^. Measurements included cortical tissue mineral density (Ct.TMD, mgHA/cm^3^), cortical bone area (Ct.Ar, mm^2^), polar and minimum moment of inertia (pMOI, I_min_, mm^4^), medullary area (Ma.Area, mm^2^), total cross-sectional area (bone + medullary area) (Tt.Ar, mm^2^), and bone area fraction (Ct.Ar/Tt.Ar, %).

### 2.6 Femoral whole-bone mechanical and material properties

Left femurs were frozen and thawed once before three-point bending. The test was performed to failure at a rate of 5 mm/min on a custom fixture with an 8 mm span, such that femurs were loaded with the posterior side facing down (1 kN load cell, Instron 5543). Femurs were hydrated before testing using PBS. Using load-displacement data and the I_min_ and C_min_ values from microCT, modulus, yield strength, maximum strength, and toughness were calculated using standard equations for the mouse femur ^(37)^.

Right femurs were assessed for notched fracture toughness following methods in alignment with prior description ^(38)^. Thawed femurs were hydrated with PBS and notched on the posterior side to a target of 1/3^rd^ of the anterior-posterior width on the posterior surface using a custom precision saw. Notched femurs were then tested with the notched side down in three-point bending at a rate of 0.001 mm/sec using an 8 mm span until failure. Fractured femurs were dried overnight at room temperature and imaged using field emission scanning electron microscopy (Zeiss SUPRA 55VP) in variable pressure mode (VPSE, 20 Pa, 15 kV) and analyzed using a custom MATLAB code to quantify cortical geometry and the initial notch angle. Fracture toughness (K_c_) was calculated from the initial notch angle and the maximum load ^(38)^.

To assess how differences in bone strength associate with metabolic profiles, we defined high- and low-strength groups for additional comparisons. Males demonstrated higher variability of strength than females. ‘Higher strength’ and ‘lower strength’ groups were defined by a threshold of ± 7% away from the mean strength for males. For females, high strength bones were those > 5% greater than the female group mean. Low strength bones were not selected for females since there was not a cluster of distinctly low strength bones for this sex.

### 2.7 Statistical analysis

The effect of sex on all bone outcomes except metabolomic measures was tested using two-sample t tests (Minitab, v.19). Data were checked for normality. Non-normal data were evaluated using the nonparametric Mann Whitney test. Significance was set *a priori* to < 0.05.

MetaboAnalyst was utilized to assess metabolomic data. Using standard procedures^(39,40)^, raw data were log transformed, standardized, and auto-scaled (mean centered divided by standard deviation per variable) prior to analysis. Analyses included hierarchical cluster analysis (HCA), principal component analysis (PCA), partial least squares-discriminant analysis (PLS-DA), volcano plot analysis, t-test, and fold change.

A concise overview of the statistical methods used in this study are discussed in detail in supplementary material. In brief, HCA and PCA are unsupervised multivariate statistical analyses that assess differences in metabolomic profiles between experimental groups. HCA identifies sub-groups of samples and determines differences between groups. PCA is an unsupervised technique that finds components (PCs) of the dataset that align with the overall variation to examine the underlying structure of the data. PLS-DA is a supervised analysis that partitions variation within the data base on *a priori* knowledge of experimental groups. PLS-DA further calculates a variable importance in projection (VIP) score to quantify how much each metabolite feature contributes to discriminating between cohorts. Taken together, these three approaches, HCA, PCA, and PLS-DA, provide a global view of the thousands of metabolites comprising the metabolome. This view describes how experimental groups are both similar and different. These complementary analyses provide insight into interactions occurring at the metabolic level.

Finally, volcano plot and fold change analyses are utilized to identify differentially regulated metabolite features between two groups. Employing these two tests allows identification of metabolites that differ in intensity between groups. These differentially expressed metabolites are then subjected to pathway analysis. Pathways are determined using MetaboAnalyst’s MS Peaks to Pathways feature with the *mummichog* algorithm. This allows for metabolite compounds identified from the statistical tests described to be associated with biological pathways and for prediction of networks of functional cellular activity. Significance was defined *a priori* as 0.05. In the case of multiple comparisons, p-values were corrected for false discovery rate (FDR) to maintain family-wise error at 0.05.

## 3. Results

### 3.1 The metabolome is distinct for whole bone, isolated cortical bone, and bone marrow

We first tested whether metabolomic profiles are distinct for whole bone (i.e., cortical and trabecular bone and marrow), isolated cortical bone, bone marrow from female mice (Supplementary Table 1). Unsupervised (HCA and PCA) and supervised (PLS-DA) multivariate statistical analyses were utilized to compare the metabolome of these three tissue groups. The global metabolomic profiles of bone marrow, isolated cortical bone tissue, and whole bone are substantially different (Figure 2A-C). HCA showed perfect clustering of samples within their respective groups (Figure 2A). PCA and PLS-DA also showed clear separation between the metabolomes of the three groups (Figure 2B, C). There were 2,178 metabolite features that were significantly different between tissues groups using ANOVA with FDR-corrected p_FDR_ < 0.05. Further, 1,120 metabolite features were detected that differed between the three experimental groups with p-value less than 0.0001 (Figure 2D). In depth analysis revealed metabolite features unique to isolated cortical bone and bone marrow. Unsupervised clustering of the median intensity for each metabolite within cohort summarizes the major differences between cohorts (Figure 2E).

**Figure 2.**
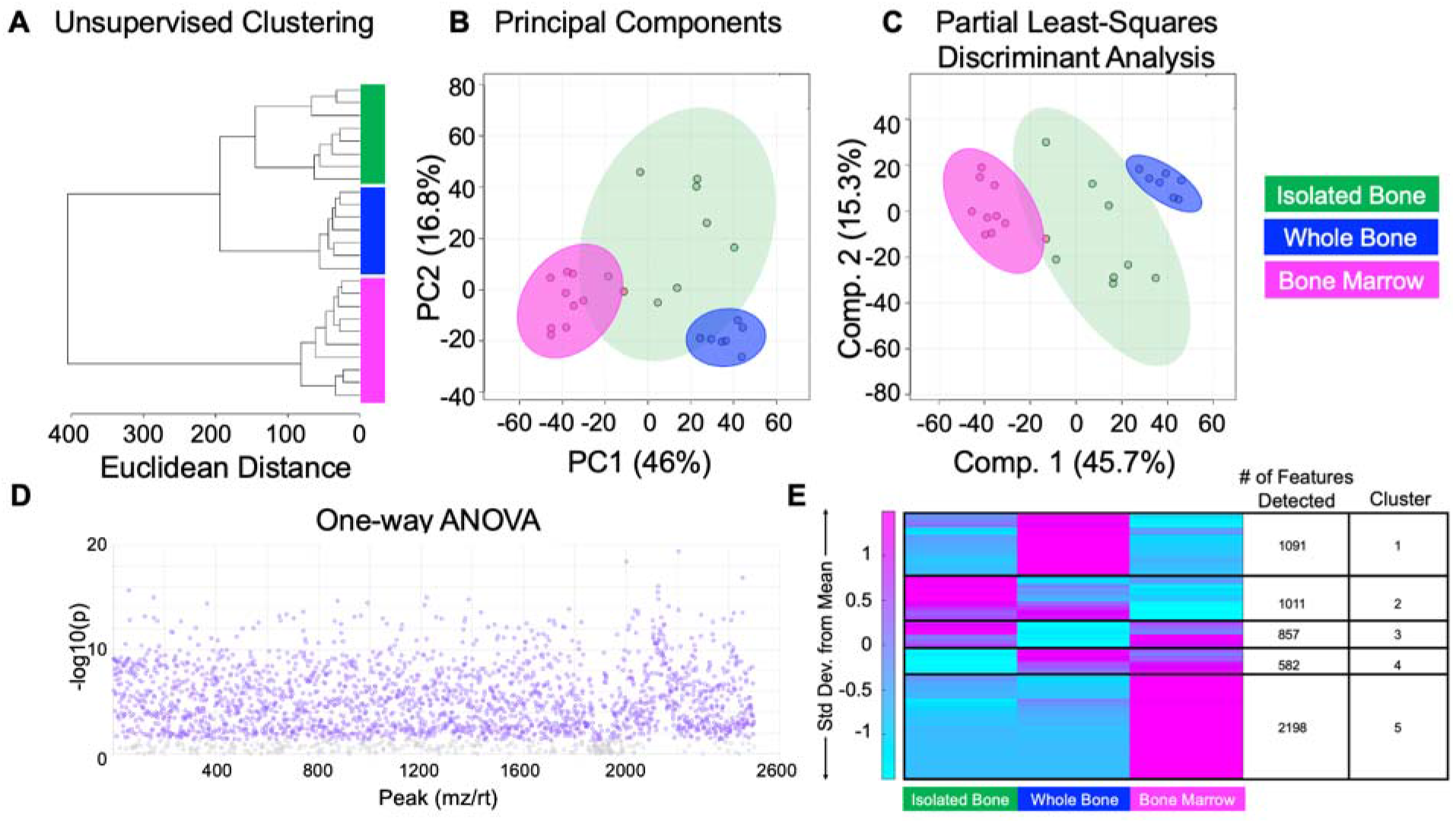
The metabolomes of whole bone, isolated cortical bone, and bone marrow are distinct. A total of 2,764 metabolite features were detected across all experimental groups including isolated bone, whole bone, and bone marrow. Features were analyzed using unsupervised (HCA and PCA) and supervised (PLS-DA) statistical methods. (A) Unsupervised HCA visualized by a dendrogram reveals that the metabolome individual tissues cluster together and are distinct from each other. (B) PCA, shown as a scatterplot, displays minimal overlap of clusters. The x axis shows PC1 which accounts for 46% of the variation in the dataset and the y axis shows PC2 which accounts for 16.8% of variation. (C) Supervised PLS-DA further displays clear separation of groups and further supports the notion that the metabolome of various tissues including isolated bone, whole bone, and bone marrow are unique. Component 1 and 2 combined accounts for 63% of variation within the dataset. The colors in A-C correspond to experimental groups: green – isolated bone, blue – whole bone, pink – bone marrow. (D) ANOVA analysis identifies over 2,000 statistically significant metabolite that are differentially regulated across tissues. (E) Heatmap analysis reveals that metabolic phenotypes greatly differ between isolated bone, whole bone, and bone marrow. Median intensities were clustered into 3 respective groups via MATLAB to visualize metabolic differences and phenotypes for tissues of interest.

Pathway analysis revealed metabolic pathways unique to bone marrow included steroid hormone biosynthesis, amino acid metabolism (phenylalanine, tyrosine, tryptophan, lysine), vitamin metabolism (biotin metabolism), and purine metabolism. Heatmap analysis identified that metabolic pathways common to cortical bone and whole bone included amino acid metabolism (cysteine, methionine, histidine, alanine), lipid metabolism (fatty acid elongation, glycosphingolipid metabolism, GPI-anchor biosynthesis, linoleic acid metabolism), electron-transport metabolism (ubiquinone metabolism), and the pentose phosphate pathway (PPP) (Table 1). Pathways unique to cortical bone include mannose type-O glycan biosynthesis, linoleic metabolism, glycosphingolipid biosynthesis, and quinone biosynthesis (ubiquinone, terpenoid-quinone) (Table 1).

**Table 1.** A) Distinct metabolic pathways associated with humerus-derived isolated cortical bone, whole bone, and bone marrow from male and female C57Bl/6 mice. B) Metabolic pathways distinct to females and males for the isolated cortical humerus. C) Metabolic pathways distinct to high strength males, high strength females, and low strength males. Pathways listed have a false discovery rate-corrected significance level < 0.05.

| **A) Musculoskeletal tissue associated pathways** |  |  |  |
| --- | --- | --- | --- |
| **Metabolic Pathway** | **Pathway Total** | **Hits** | **Tissue** |
| Glycosaminoglycan degradation | 21 | 13 | Whole Bone |
| Pentose phosphate pathway | 22 | 5 | Whole Bone |
| Cysteine and methionine metabolism | 33 | 6 | Whole Bone |
| Fatty acid elongation | 30 | 4 | Whole Bone |
| Histidine metabolism | 16 | 7 | Whole Bone |
| Ubiquinone and other terpenoid-quinone biosynthesis | 9 | 6 | Isolated Cortical Bone |
| Mannose type O-glycan biosynthesis | 16 | 6 | Isolated Cortical Bone |
| Linoleic acid metabolism | 4 | 3 | Isolated Cortical Bone |
| Glycosphingolipid biosynthesis | 21 | 3 | Isolated Cortical Bone |
| Beta-Alanine metabolism | 21 | 9 | Whole + Isolated Bone |
| Glycosylphosphatidylinositol (GPI)-anchor biosynthesis | 11 | 6 | Whole + Isolated Bone |
| Steroid hormone biosynthesis | 77 | 61 | Bone Marrow |
| Phenylalanine, tyrosine and tryptophan biosynthesis | 4 | 2 | Bone Marrow |
| Lysine degradation | 19 | 10 | Bone Marrow |
| Phenylalanine metabolism | 12 | 4 | Bone Marrow |
| Biotin metabolism | 4 | 4 | Bone Marrow |
| Purine metabolism | 66 | 20 | Bone Marrow |
| **B) Sexually dimorphic pathways** |  |  |  |
| Alanine, aspartate and glutamate metabolism | 28 | 4 | Male |
| Arginine and proline metabolism | 37 | 2 | Male |
| Cysteine and methionine metabolism | 33 | 4 | Male |
| Purine metabolism | 66 | 7 | Male |
| Glycolysis / Gluconeogenesis | 23 | 3 | Male |
| Citrate cycle (TCA cycle) | 16 | 3 | Male |
| Sphingolipid metabolism | 9 | 2 | Female |
| Glycerophospholipid metabolism | 13 | 1 | Female |
| GPI-anchor biosynthesis | 11 | 3 | Female |
| Fatty acid biosynthesis | 10 | 2 | Female |
| **C) Strength associated pathways** |  |  |  |
| Purine metabolism | 66 | 10 | High Strength Males |
| Porphyrin metabolism | 27 | 6 | High Strength Males |
| Pyrimidine metabolism | 39 | 7 | High Strength Males |
| Tryptophan metabolism | 41 | 7 | High Strength Males |
| beta-Alanine metabolism | 21 | 3 | High Strength Males |
| Aminoacyl-tRNA biosynthesis | 22 | 3 | High Strength Males |
| Terpenoid backbone biosynthesis | 15 | 4 | High Strength Females |
| Tryptophan metabolism | 41 | 7 | High Strength Females |
| Porphyrin metabolism | 27 | 6 | High Strength Females |
| Pentose phosphate pathway | 22 | 2 | Low Strength Males |
| Pantothenate and CoA biosynthesis | 17 | 3 | Low Strength Males |
| Phosphatidylinositol signaling system | 17 | 3 | Low Strength Males |

### 3.2 The cortical bone metabolome differs by sex

We then compared the isolated cortical bone metabolome for female and male mice. A total of 2,129 distinct metabolite features were detected. These features were further analyzed (Figure 3). HCA showed that males and females cluster separately except for one male mouse (Figure 3A). We then utilized PCA to analyze the overall variation in the dataset between males and females. Principal component 1 accounted for 24% of the variation in the dataset (i.e., if the metabolomes were the same, principal components would account for 1/(2,129 metabolite features x 20 samples) = 0.00002% of the variation in the dataset).This result strongly suggests that the metabolome for isolated cortical bone from males and females is different (Figure 3B).

**Figure 3.**
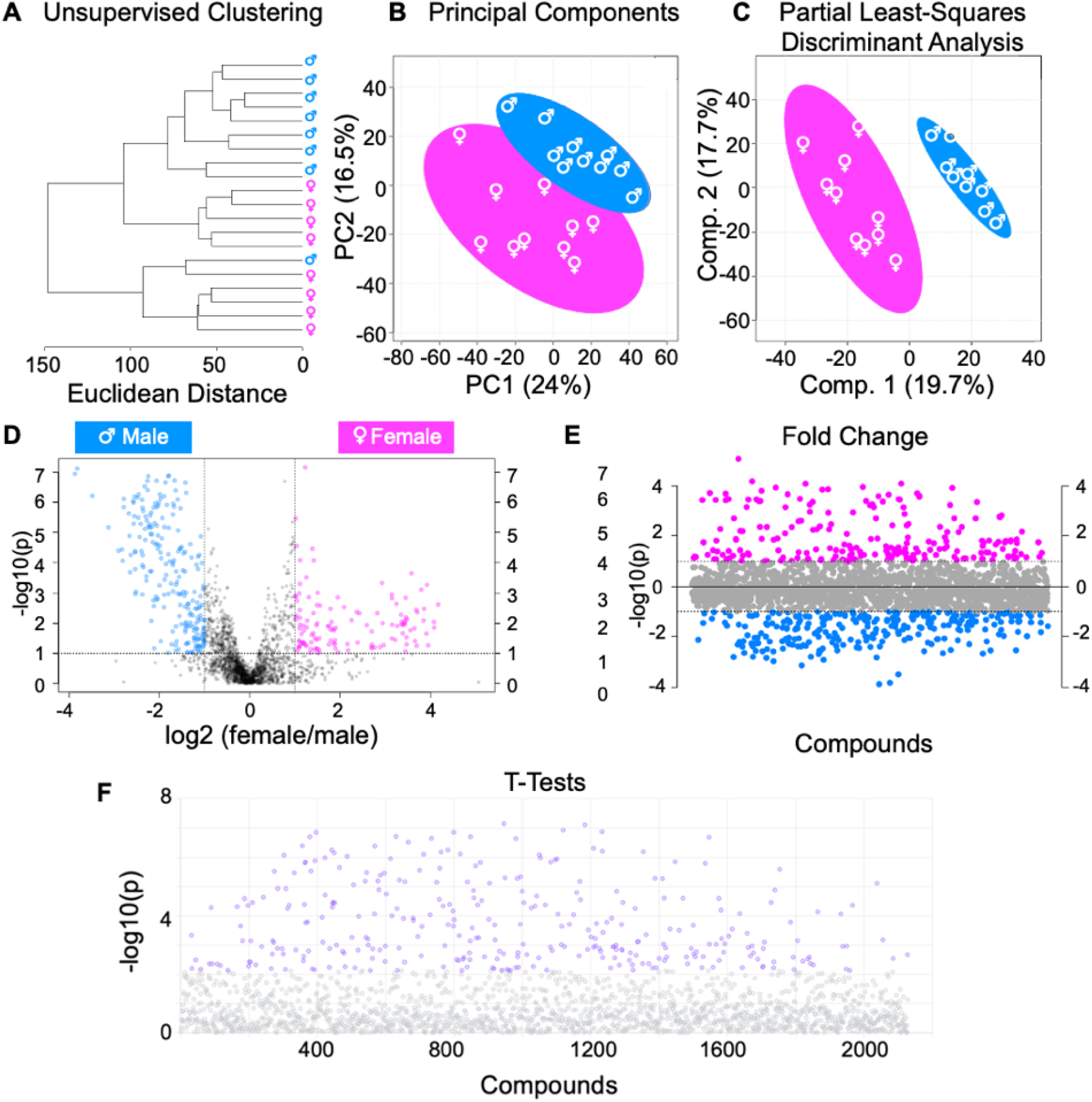
Metabolomic profiles of isolated bone show sexual dimorphism. A total of 2,129 metabolite features were detected and analyzed by both unsupervised hierarchical clustering (HCA) and principal component analysis (PCA) and supervised partial least-squares discriminant analysis (PLS-DA). (A) Unsupervised HCA visualized by a dendrogram displays distinguished clusters between male and female mice, except for one male mouse clustering amongst females. (B) PCA, like HCA, displays distinguished clusters with minimal overlap of males and females. PCA is shown as a scatterplot with the first two PC on the x and y axes. The x axis shows PC1 which accounts for 24% of the variation in the dataset. PC2 is o the y-axis and accounts for 16.5% of the variation in the dataset. (C) Supervised PLS-DA finds clear separation between the metabolomes of male and female mice. PLS-DA is also shown as a scatterplot of the top two components, with component 1 accounting for 19.7% and component 2 accounting for 17.7% of variation within the dataset. The colors in A-C correspond to sample cohorts: blue – control males, pink – control females. (D) Volcano plot analysis displays differentially regulated metabolite features that were distinguished between cohorts using both fold change and false discovery rate (FDR) adjusted p-value. When comparing males and females, 126 metabolite features in the upper right quadrant had a fold change > 3 and a p-value < 0.05 and these features were significantly higher in control female mice. Similarly, 225 features in the upper left quadrant were associated with control male mice. (E) Fold change analysis was utilized to further examine the differences in metabolomes of female and male mice. 211 metabolite features (p-value < 0.05) with a positive FC correspond to female mice, and 267 metabolite features with a negative FC correspond to male mice. (F) T-test analysis identified 318 significant metabolites (p-value < 0.05).

We used PLS-DA to further examine the variation between the male and female cortical bone metabolome. This supervised analysis produced a complete separation between the metabolomes of male and female mice (Figure 3C). VIP scores from PLS-DA analyses were used to identify specific metabolite features that contribute the most to differences between male and females. Of the top 300 metabolite features that contributed to the distinction between groups, the majority (75%) were upregulated in male mice. Significant pathways that contributed to the separation of groups and are primarily associated with male isolated cortical bone included amino acid metabolism (cysteine, methionine, arginine, proline) and central energy metabolism (glycolysis, TCA cycle).

We then utilized T-test, fold change, and volcano plot analysis to identify specific metabolite features that differ between male and female cortical bone (Figure 3). Based on *a priori* significant level of 0.05, volcano plot analysis revealed 126 metabolite features were upregulated amongst female mice and 225 were upregulated amongst male mice (Figure 3D). Fold change analysis yielded 487 metabolite features that were differentially regulated between groups (Figure 3E). Student’s t-test identified 318 metabolite features (Figure 3F).

Numerous metabolic pathways were found for male and female cortical bone (Table 1). The main metabolic theme upregulated for males compared with females was amino acid metabolism. Upregulated metabolites included cysteine, methionine, alanine, aspartate, and glutamate. Central energy metabolism, including glycolysis, the TCA cycle, and purine metabolism, were also upregulated in males. The main metabolic theme upregulated for females was lipid metabolism. This includes sphingolipid metabolism, GPI-anchor biosynthesis, glycerophospholipid metabolism, and the fatty acid pathways of biosynthesis, degradation, and elongation.

### 3.3 Females have increased cortical bone adiposity and increased serum biomarkers of resorption

Since lipid metabolism was prominent in the female bone metabolome, we sought to investigate if metabolic upregulation of lipid metabolism corresponds to an upregulation in adiposity in females compared to males. Adipocyte count and number of adipocytes per marrow cavity area were 81% (p < 0.0001) and 83% (p < 0.0001) higher in females compared to males (Figure 4). Thus, metabolic and histological data agree that females had both increased bone marrow adiposity as well as an increased metabolic signature of lipid metabolism.

**Figure 4.**
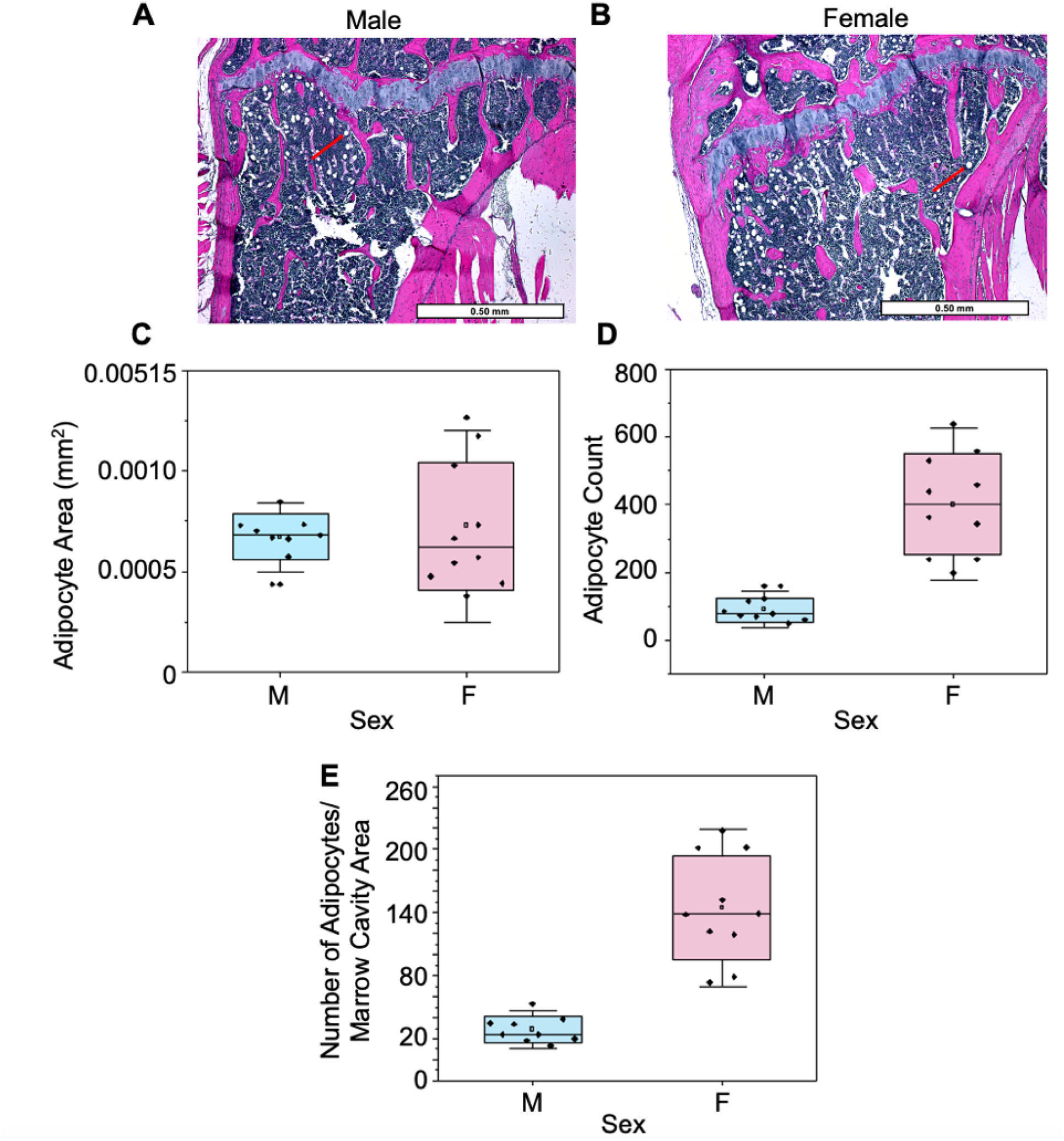
Bone marrow adiposity exhibits sexual dimorphic behavior. (A-B) Male (right) and female (left) H&E-stained proximal tibia imaged at 4x. Red arrows point to adipocytes. Measurements calculated include (C) adipocyte size (mm^2^), (D) adipocyte count and (E) number of adipocytes per marrow cavity area in male and female mice.

We also assessed serum biomarkers to estimate global bone resorption and formation activities._CTX1, a biomarker of global bone resorption, was higher in females compared to males (+51.4%, p=0.011). P1NP, a biomarker of global bone formation, was lower in females compared to males (−22.8%, p=0.027). The ratio of CTX1 to P1NP was higher in females compared to males (+ 87.1%, p = 0.001) (Figure 5, Supplemental Table 1).

**Figure 5.**
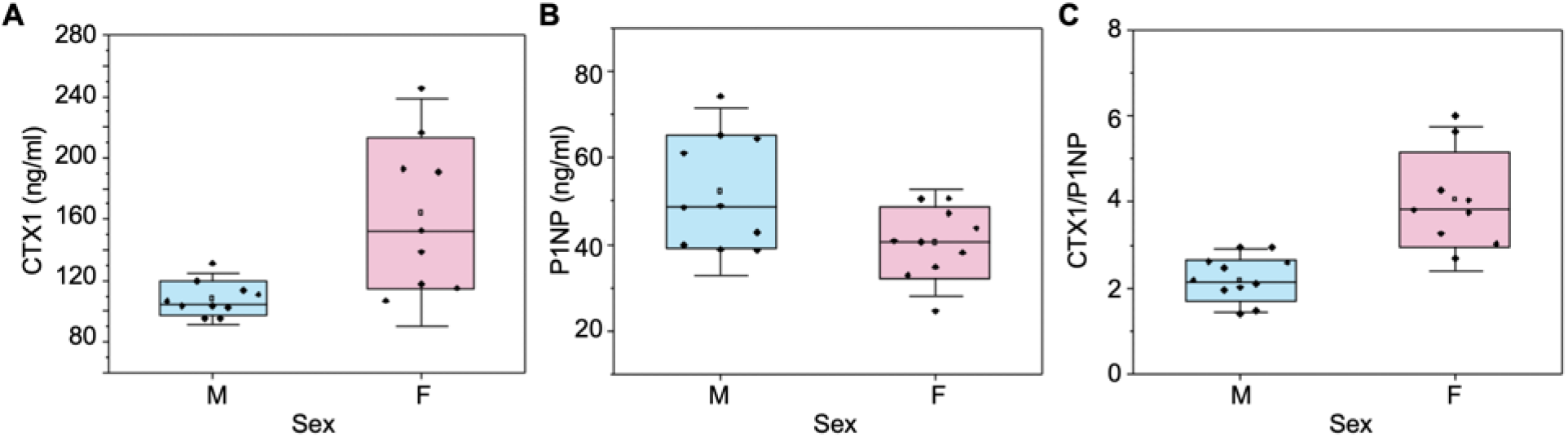
Global serum biomarkers, P1NP and CTX1, differ in concentration between male and female mice. (A) CTX1, a biomarker of bone resorption, concentration was higher in female mice compared to males. (B) P1NP, a biomarker of bone formation, concentration was higher in males compared to females. (C) The ratio of CTX1/P1NP is a measure of net bone resorption compared with formation. This measure was higher for females (p<0.05).

### 3.4 Femur trabecular and cortical microarchitecture differ by sex

Sex differences were observed in the trabecular and cortical microarchitecture of the distal femur metaphysis (Supplementary Table 1). Males had greater bone volume fraction (+64.1%, p < 0.001), trabecular tissue mineral density (+41.2%, p < 0.001), connectivity density (+66.14%, p < 0.0001), trabecular thickness (+12.71, p = 0.001), trabecular number (+29%, p < 0.0001), cortical area (+ 12.2%, p = 0.003), minimum moment of inertia (+ 39.8%, p < 0.0001), polar moment of inertia (+ 38.2%, p < 0.0001), total area (+ 26.8%, p < 0.0001), and medullary area (+ 36.8%, p < 0.0001). Females had higher structural model index (+50%, p < 0.003), bone surface to bone volume (+24%, p < 0.0001), trabecular spacing (+31.5%, p < 0.0001), cortex tissue mineral density (+4%, p < 0.0001), cortical thickness (+9.2%, p = 0.001), and cortex area/total area (+17%, p < 0.0001) (Supplementary Table 1).

### 3.5 Femur material properties differ with sex

Whole-bone material properties differed by sex (Table 2). Females had higher modulus (+25%, p < 0.0001), ultimate stress (+ 14.4%, p < 0.0001), and yield stress (+18.5%, p = 0.001) than males. No sex differences were found for either notched fracture toughness (Kc) or toughness from three-point bending (e.g., area under stress-strain curve for three-point bending of unnotched femurs).

**Table 2:**
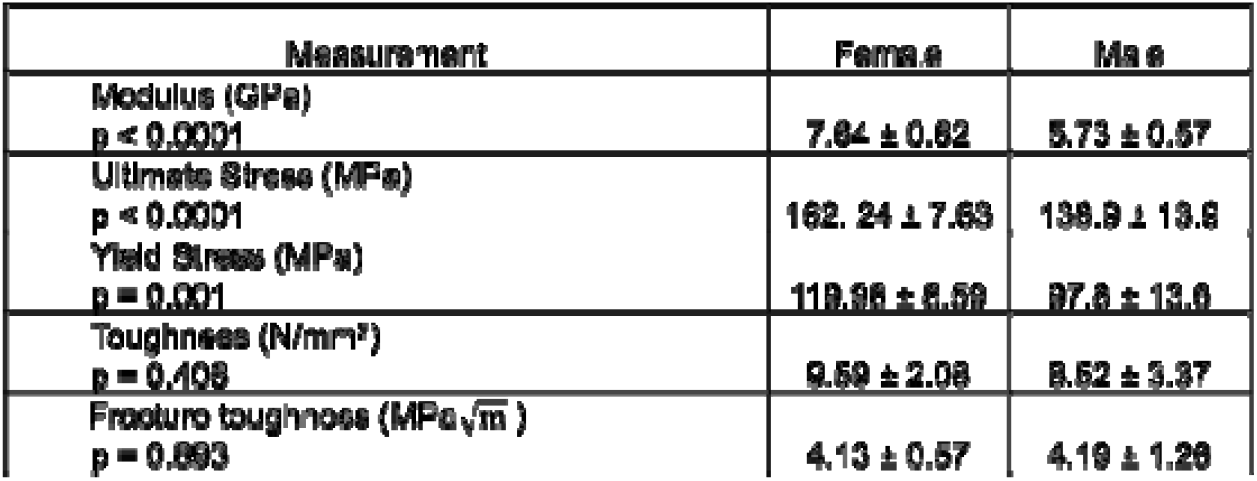
Femur material properties from 3-point bending. Data are presented as mean ± standard deviation from the mean.

### 3.6 Sex differences in the cortical bone metabolome are associated with sex differences in bone strength

Females in this study had higher femur strength than males. While females did not have much spread in strength (coefficient of variation: 4.7%), males showed more variation (coefficient of variation: 7.6%). Thus, we evaluated whether the cortical humerus metabolome differed for males with the highest strength and highest lowest strength femurs. We also evaluated whether the cortical humerus metabolome differed between males and femurs with the highest strength femurs of their sex.

The highest strength (n=4) and lowest strength femurs (n=4) from males and only the strongest female femurs (n=3) were grouped (males = ± 7%, females = + 5% difference from the mean values for each sex). There was not a distinct lower-strength group for females, so only higher strength femurs were grouped. We employed HCA, PCA, and PLS-DA to assess if metabolomic profiles were associated with each strength group (Figure 6). High strength females, high strength males, and low strength males had partial separation in metabolomic profiles, as seen from HCA (Figure 6A). PCA found differences between samples within their respective cohorts. PC2 and PC3 were analyzed and together accounted for almost 30% of the overall variation in the dataset (Figure 6B, C). PLS-DA showed that each of the three strength groups are metabolically distinct from each other (Figure 6D).

**Figure 6.**
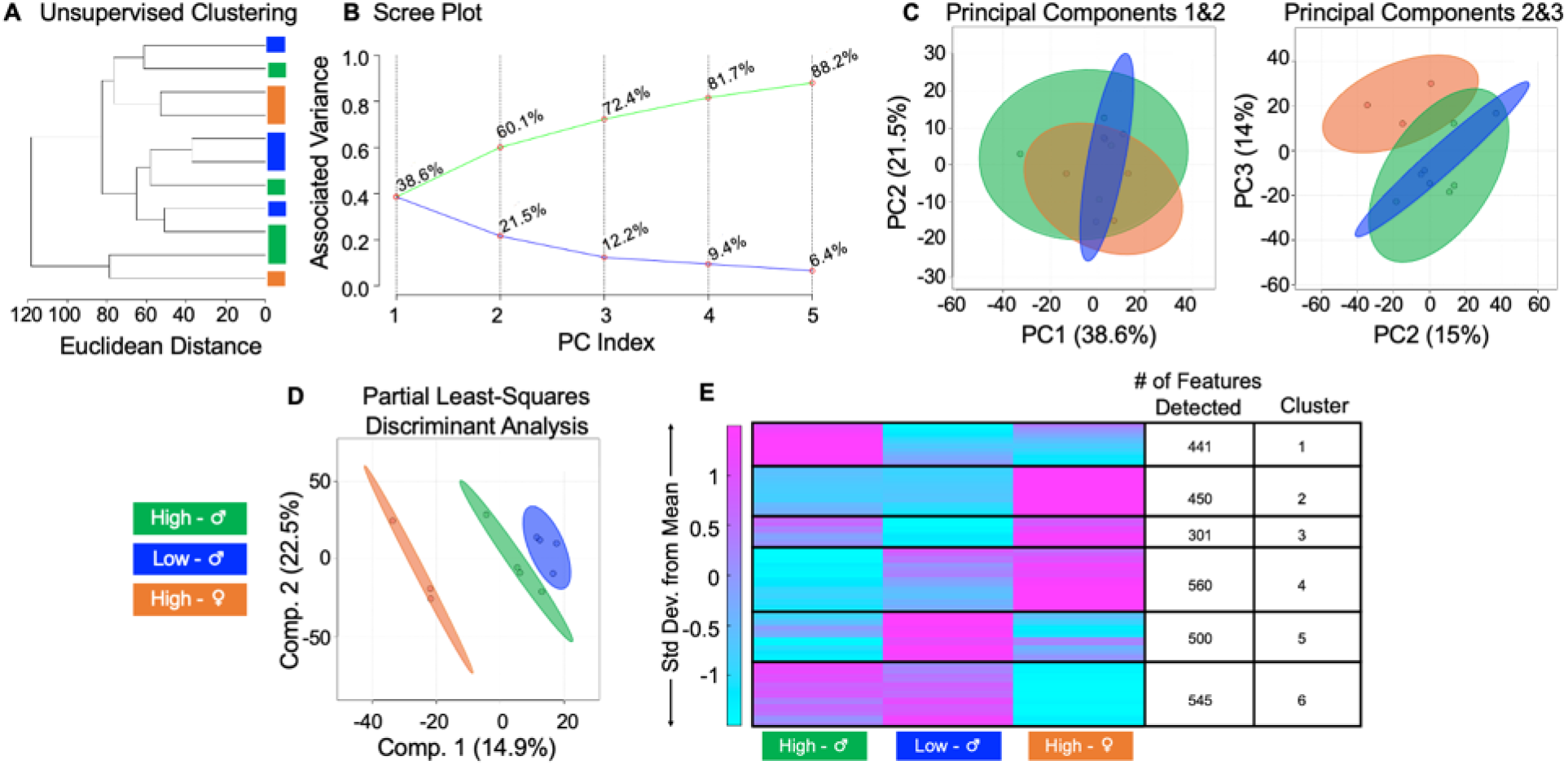
Differences in whole bone strength correspond to metabolic differences in male and female mice. An untargeted metabolomic approach was utilized to generate metabolomic profiles based on whole bone strength. (A) HCA, visualized by a dendrogram, displays that the metabolome of bone differing by strength somewhat differs. (B) Scree plot reveals how much variation is being captured in each principal component from the data. (C) PCA analysis, captured by PC1 and PC2 show moderate overlap of groups. Together, PC1 and PC2 account for approximately 60% of the variation in the dataset. PCA analysis captured by PC2 and PC3 show improved separation compared to B. (D) PLS-DA analysis displays complete separation of groups. Similarly, component 1 and 2 combined account for 30% of the variation in the dataset. (E) Median intensity heatmap analysis shows that the metabolome of stronger and weaker bones differs from each other, and when accounting for sex. Median intensities were clustered into 3 respective groups via MATLAB to visualize and identify metabolic pathways and features that differ between groups. The colors in A-E correspond to sample cohorts: green – high strength males, blue – low strength males, orange – high strength females.

Unique pathways and features for each strength group were identified using HCA and heatmaps (Figure 6E). The data suggest that strong female femurs are associated with terpenoid backbone biosynthesis compared to high- and low-strength males (Table 1). Metabolic pathways shared by males and females with stronger femurs included tryptophan and porphyrin metabolism. Metabolic themes unique to strong male femurs included purine and pyrimidine metabolism, beta-alanine metabolism, and aminoacyl tRNA biosynthesis. The metabolome of males with weaker femurs included the PPP, pantothenate and CoA biosynthesis, and phosphatidylinositol signaling system.

Our data show that differences in bone strength are associated with substantial differences in isolated bone metabolomic profiles. In this study, bone strength had moderate to strong correlations with measures of bone turnover. For pooled female and male data (all mice in the study), ultimate strength was negatively correlated with P1NP (Spearman’s ρ = −0.486, p-value = 0.03) and was positively correlated with CTX1 (Spearman’s ρ= 0.527, p-value=0.02) and CTX1/P1NP (Spearman’s ρ= 0.707, p-value=0.01) (Supplementary Figure 2). For males, strength was not correlated with CTX1 (p= 0.036, ρ > 0.05) and may be moderately correlated with P1NP (ρ= −0.418, ρ > 0.05) and CTX1/P1NP (ρ = 0.588, p = 0.07). For females, strength was not related to P1NP (ρ = −0.018, p > 0.05) and may be weakly to moderately related to CTX1 (ρ= – 0.233, p > 0.05) and CTX1/P1NP (ρ= −0.4, p > 0.05). Thus, some of the differences between male and female strength groups, and between higher and lower strength males, may reflect differences in bone turnover.

## 4. Discussion

This study evaluated metabolomic profiles of murine humeri to provide new insight into how molecular processes associated with different bone tissues (e.g., cortical bone, bone marrow, whole bone) and with female and male sex. Metabolomic profiling has been used in other areas of orthopedic research ^(39–55)^. For example, metabolomic assessment revealed that synovial fluid phenotypes reflect cartilage morphological changes and disease progression ^(40,42–44,46,53)^. Metabolomic profiling has also contributed to biomarker and drug targets for diseases like osteoarthritis^(42,46)^ and rheumatoid arthritis ^(40,47,56)^. Metabolomics is an attractive technique for cortical bone tissue because it can identify differences in metabolites produced by the same bone cells that influence cortical bone quality. However, metabolomic analyses are thus far infrequently applied to cortical bone tissue and key questions remain unanswered about the dependence of metabolomic profiles on bone tissue type and sex.

### 4.1 The metabolomes of cortical bone, bone marrow, and whole bone are distinct

We found that metabolomic profiles of isolated cortical bone, bone marrow, and whole bone of the humerus are distinct. Pathways upregulated in bone marrow included steroid metabolism, vitamin metabolism, and purine metabolism. A common upregulated pathway shared by all three tissues was amino acid metabolism. However, different amino acids were upregulated in each tissue. Phenylalanine, tryptophan, tyrosine, and lysine metabolism were upregulated in bone marrow. These aromatic amino acids are commonly grouped together because of their aromaticity and because they feed into, or can be the products of, glycolysis and the TCA cycle ^(57)^. These findings are consistent with the paradigm of bone marrow as an active endocrine organ, with many cells that serve as a long-term energy reserve ^(58)^. It is important to note that bone marrow is composed of numerous cell types and is influenced by sex, age, and endocrine factors ^(33)^. Therefore, metabolic signatures of flushed humeri-derived bone marrow may be driven by many unique factors.

Pathways upregulated in isolated cortical bone and whole bone, but not marrow, included cysteine, methionine, histidine, and beta-alanine metabolism. Methionine is an essential amino acid that plays a large role in several metabolic processes including nucleotide synthesis. Methionine restriction prolongs lifespan in mice. However, methionine restriction also has negative impacts on bone density, overall bone structure, and the innate immune system ^(59)^. Through the conversion of methionine to homocysteine, a common downstream product of methionine metabolism is cysteine ^(60)^. Cysteine, along with other glucogenic amino acids, can be used as an energy source by conversion into glucose. In postmenopausal women, low cysteine has been associated with low BMD and increased bone turnover ^(61)^. Dysregulation of cysteine, and upstream methionine, can lead to bone fracture and loss ^(62)^. Elevated levels of histidine in orthopedic tissues, including synovial fluid and cartilage, is found both in response to mechanical stimulation and osteoarthritis ^(63,64)^. Beta-alanine was elevated in synovial fluid, serum, and subchondral bone in osteoarthritis animal models ^(44,52,65)^. Pathways unique to cortical bone include mannose type-O glycan biosynthesis, linoleic metabolism, glycosphingolipid biosynthesis, and quinone biosynthesis (ubiquinone, terpenoid-quinone). Isolated cortical bone and whole bone differ not only from marrow flushing but also because whole bone includes trabecular bone. Therefore, pathways that are different between cortical bone and whole bone that do not align with those of bone marrow may also reflect differences apparent between cortical and trabecular compartments.

### 4.2 Bone is sexually dimorphic across many length-scales, including the metabolome

Our results build on previous work that mouse bone properties are sexually dimorphic ^(1–16)^. We observed that male mice have greater femur trabecular bone microarchitecture and have higher serum biomarkers of bone formation. Conversely, females had stronger and stiffer femurs, greater cortical thickness and tissue mineral density, and increased serum biomarkers of bone resorption. This is the first work, to our knowledge, to assess sex differences in the cortical bone metabolome.

Males had elevated P1NP, lower CTX1, and upregulated amino acid metabolism. Metabolic pathways previously associated with P1NP include energy metabolism (e.g., TCA cycle), amino acid metabolism, and pyrimidine metabolism ^(66)^. Energy and amino acid metabolism were also upregulated in male isolated cortical bone. Upregulated amino acids included alanine, aspartate, glutamate, cysteine, methionine, proline, and arginine. Alanine is generated from pyruvate and is therefore closely linked to glycolysis and the TCA cycle. Alanine has been shown to aid in the recycling of carbon backbones in skeletal muscle and the liver ^(67)^. Glutamate is an excitatory neurotransmitter, and there is a growing body of evidence of a regulatory role for glutamate in osteoblast and osteoclast differentiation, as well as bone homeostasis ^(68–72)^. Our metabolomic and serum biomarker analyses suggest that the elevated P1NP concentration in males is associated with the upregulation of energy and amino acid metabolism (Figure 5–6, Table 1, Supplementary Table 2).

Female mice had greater bone marrow adiposity and upregulated lipid metabolism (Table 1). The metabolism of lipids is essential for energy regulation, membrane dynamics, and signaling ^(73)^. Fatty acids and other lipids are transported systemically via chylomicrons to be cleared by the liver and bone. After the liver, the femur diaphysis is the second most active organ for chylomicron reuptake, supporting the idea that normal osteoblast proliferation is fueled by serum lipoproteins ^(74,75)^. This is consistent with the canonical role of bone’s importance in fatty acid clearance for energy purposes ^(76,77)^. When lipids are limited, osteogenesis is negatively influenced because osteoblasts depend on fatty acid oxidation ^(78)^. Therefore, greater adiposity and upregulation of lipid metabolism in females may demonstrate usage of fat as a substrate. An upregulation of fatty acid metabolism for females may demonstrate usage of fat as a substrate. It is possible that this increased lipid metabolism for females has a functional role for bone resorption. We observed that females had greater biomarkers of bone resorption (CTX1 and CTX/P1NP) than males. In a study investigating healthy young adult serum using metabolomics and biomarker analysis CTX1 was positively correlated with lipid metabolism (fatty acid biosynthesis), beta oxidation, and carbohydrate metabolism in humans ^(66)^. Previous studies found that osteoclasts’ high energy demand can be fueled by lipids ^(66,79–81)^, but it is uncertain if osteoblasts and osteocytes are fueled by lipids in the same way. Our study utilized marrow-flushed cortical bone. It is possible that metabolic products of osteoclasts may still be present within cortical tissue.

### 4.3 Bone metabolism differs in association with bone strength and bone turnover

Because bone turnover strongly influences bone strength ^(82)^ and requires cellular energy production and utilization ^(23,24)^, we hypothesized that the cortical bone metabolome would differ between groups of different bone strength. The cortical bone metabolomes of high strength males had upregulated nucleotide metabolism, while high strength females had upregulated terpenoid backbone biosynthesis and low strength males had an upregulation of the PPP (Table 1).

Both high strength male and female mice had upregulated tryptophan metabolism. Tryptophan is a precursor to serotonin, melatonin, and kynurenine ^(83–85)^. In humans, downstream metabolites of tryptophan metabolism, like kynurenic acid, have been shown to influence bone remodeling by inhibiting glutamate receptors ^(70,84,86)^. Other mechanisms of tryptophan prominently affecting bone remodeling is through stimulating proliferation and differentiation of osteoblasts and bone marrow mesenchymal stem cells ^(87)^. In both humans and mice, tryptophan has been associated with osteoclast activity and was positively associated with CTX1 levels ^(66)^. In our study, there was a moderate correlation between bone strength and CTX1/P1NP for males. Females also had higher bone strength and higher CTX1/P1NP than all males. Therefore, the detection of upregulated tryptophan metabolism for high strength males and females is consistent with the increased global bone resorption observed for the same mice.

High strength males had upregulated nucleotide metabolism (i.e., purine and pyrimidine metabolism) compared to high strength females and low strength males. In low strength males, the PPP was upregulated. Both purine and pyrimidine metabolism are derived from the PPP. Purine and pyrimidine metabolism are important pathways for DNA and amino acid synthesis ^(88)^, which are consistent with the relatively increased bone formation relative to resorption (e.g., lower CTX1/P1NP) for low strength males. This finding may represent a higher cellular demand for nucleotides. However, additional studies are needed to determine the mechanistic details and relevance of nucleotide metabolism to bone strength.

Overall, the metabolic differences between high- and low-strength bone were different than those found between the broader groups of all female and male mice. For example, when comparing all males and females, lipid metabolism was upregulated for females and amino acid metabolism was upregulated in males. By contrast, when comparing high strength males and females, terpenoid backbone biosynthesis was upregulated in high strength females and purine and pyrimidine were upregulated in high-strength males. Our data suggest that metabolic differences between male and female strength groups likely reflect sex differences in bone turnover, since females have higher CTX1/P1NP (Supplementary Figure 2).

### 4.4 Limitations

There are important limitations to this study. First, the femur was used for mechanical testing while humeri were used for metabolomic assessments. Therefore, the metabolism and material and mechanical properties of other bones may differ. Second, marrow-flushing can leave trace amounts of marrow, although from our histological assessments we estimate that marrow remaining was very minimal (Supplementary Figure 1). Third, while female and male C57Bl/6J mice were for experiments involving bone quality characterization, only female C57Bl/6J mice were used to assess metabolic differences between whole bone, isolated bone, and bone marrow. Finally, humeri were not weighed, therefore, it is unknown if all bones had the same amount of cellular content.

### 4.5 Summary

Bone quality is sexually dimorphic for C57Bl/6 mice ^(1,2,4–7,9,10,12–14,16)^. Our data demonstrate that the cortical bone metabolome is distinct from marrow and is also sexually dimorphic. We found that males and females rely on different metabolic pools and pathways to generate ATP and meet energy demands. For example, female mice predominantly utilized lipid metabolism to meet energy demands, whereas males utilized amino acid metabolism. Males and females with higher values of bone strength had upregulated tryptophan metabolism. Since the stronger groups (e.g., high strength males vs low strength males; high strength females vs high or low strength males) had higher CTX1/P1NP, we estimate that the metabolomic signature of bone strength in our study least partially reflects differences in bone turnover. The sex differences evident in the cortical bone metabolome may help to connect sex differences in bone cell health and behavior with tissue-level differences in bone quality.

## Supporting information

Supplement File 3 Raw Metabolomics Data

## Acknowledgements

We thank the Montana State University Mass Spectrometry Facility and Dr. Katie Steward for assisting in LC-MS analysis. Funding for the Proteomics, Metabolomics and Mass Spectrometry Facility used in this publication was made possible in part by the MJ Murdock Charitable Trust, the National Institute of General Medical Sciences of the National Institutes of Health under Award Numbers P20GM103474 and S10OD28650, and the MSU Office of Research, Economic Development and Graduate Education. Additionally, assistance from Maria Jerome at the Montana State University Histology Core Facility, and Dr. Heidi Smith and Dr. Markus Dieser at the Center for Biofilm Engineering at Montana State University in histological preparation, imaging, and analysis is gratefully acknowledged. We acknowledge the Center for Advanced Orthopaedic Research for microCT analyses. Finally, we thank Maya Moody, Leah Davidson, Kenna Brown, and Priyanka Brahmachary for assisting with tissue harvests. Funding was provided by the National Science Foundation (CMMI 1554708, CMMI 2120239) and the National Institutes of Health (NIAMS R01AR073964, NIGMS P20GM103474, NIH R03AG068680).

## Author contributions

HDW performed dissections, extracted metabolites, analyzed data, and drafted the manuscript. GV performed dissections, bone flexural testing, and analyzed data. STW supplied study mice. BB assisted in analyzing data. SAM provided serum biomarker analysis materials and assisted with analysis. CMH designed experiments and analyzed data. RKJ designed experiments and analyzed data. All authors have read and revised the manuscript.

## Supplementary Materials

**Supplementary Table 1.** Serum biomarkers, P1NP and CTX1, concentrations for different experiment groups. Data are presented as mean ± standard deviation from the mean.

| Group | P1NP (ng/ml) | CTX1 (ng/ml) | CTX1/P1NP |
| --- | --- | --- | --- |
| High Strength Females | 40.79 ± 2.83 | 128.47 ± 21.03 | 3.16 ± 0.55 |
| High Strength Males | 46.69 ± 12.53 | 110.44 ± 12.30 | 2.42 ± 0.28 |
| Low Strength Males | 59.62 ± 15.10 | 108.05 ± 10.67 | 1.94 ± 0.71 |

**Supplementary Table 2.** Trabecular and cortical microarchitecture and cortical geometry from microCT. Data are presented as mean ± standard deviation from the mean. BV/TV = bone volume/total volume; BMD = bone mineral density; Conn.D = connective density; SMI = structural model index; BS/BV = bone surface to bone volume; Tb.Th = trabecular thickness; Tb.N = trabecular number; Tb.S = trabecular spacing; Ct. TMD = cortex tissue mineral density; Ct. Area = cortical area; Imin = minimum moment of inertia; Ct. Th = cortical thickness; pMOI = polar moment of inertia; Tt. Area = Total Area; Ct. Area/ Tt. Area = Cortical Area/Total Area; Ma. Area = medullary area.

| Measurement | Female | Male |
| --- | --- | --- |
| Trabecular Microarchitecture |  |  |
| BV/TV (%)<br>p < 0.0001 | 6.51 ± 1.10 | 18.12 ± 0.89 |
| BMD (mgHA/cm<sup>3</sup>)<br>p < 0.0001 | 130.82 ± 10.83 | 222.51 ± 23.33 |
| Conn.D (1/mm<sup>3</sup>)<br>p < 0.0001 | 33.82 ± 8.71 | 99.88 ± 11.30 |
| SMI<br>p < 0.0001 | 3.08 ± 0.25 | 1.54 ± 0.37 |
| BS/BV (mm<sup>2</sup>/mm<sup>3</sup>)<br>p < 0.0001 | 56.43 ± 4.85 | 42.66 ± 4.77 |
| Tb.Th (mm)<br>p = 0.001 | 0.0515 ± 0.0039 | 0.0590 ± 0.0045 |
| Tb.N (1/mm)<br>p < 0.0001 | 3.02 ± 0.25 | 4.25 ± 0.22 |
| Tb.Sp (mm)<br>p < 0.0001 | 0.33 ± 0.03 | 0.23 ± 0.01 |
| Cortical Microarchitecture |  |  |
| Ct. TMD (mgHA/cm<sup>3</sup>)<br>p < 0.0001 | 1244.00 ± 10.50 | 1198.80 ± 9.75 |
| Ct. Area (mm<sup>2</sup>)<br>p = 0.003 | 0.853 ± 0.074 | 0.971 ± 0.078 |
| Imin (mm<sup>4</sup>)<br>P < 0.0001 | 0.119 ± 0.013 | 0.198 ± 0.026 |
| Ct. Th (mm)<br>p = 0.001 | 0.197 ± 0.011 | 0.179 ± 0.009 |
| pMOI (mm<sup>4</sup>)<br>p < 0.0001 | 0.389 ± 0.061 | 0.630 ± 0.093 |
| Tt. Area (mm<sup>2</sup>)<br>p < 0.0001 | 1.75 ± 0.108 | 2.39 ± 0.182 |
| Ct. Area/Tt. Area (%)<br>p < 0.0001 | 48.69 ± 2.41 | 40.66 ± 1.81 |
| Ma. Area (mm<sup>2</sup>)<br>p < 0.0001 | 0.8970 ± 0.0612 | 1.419 ± 0.123 |

**Supplementary Table 3.** Raw metabolomics data. See attached.

**Supplementary Figure 1.**
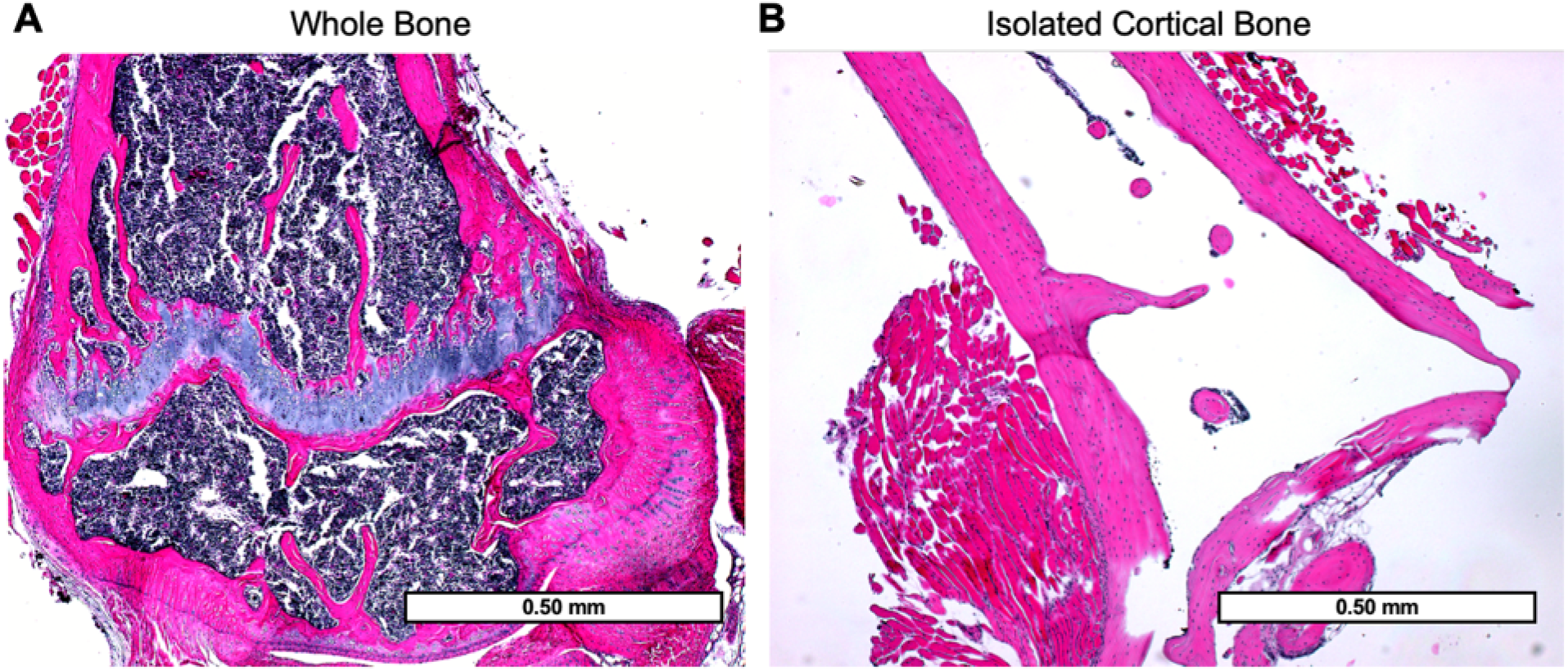
H&E stained sections to assess the adequacy of marrow flushing. (A) Whole bone and (B) marrow-flushed isolated cortical bone H&E-stained humeri imaged at 4x.

**Supplementary Figure 2.**
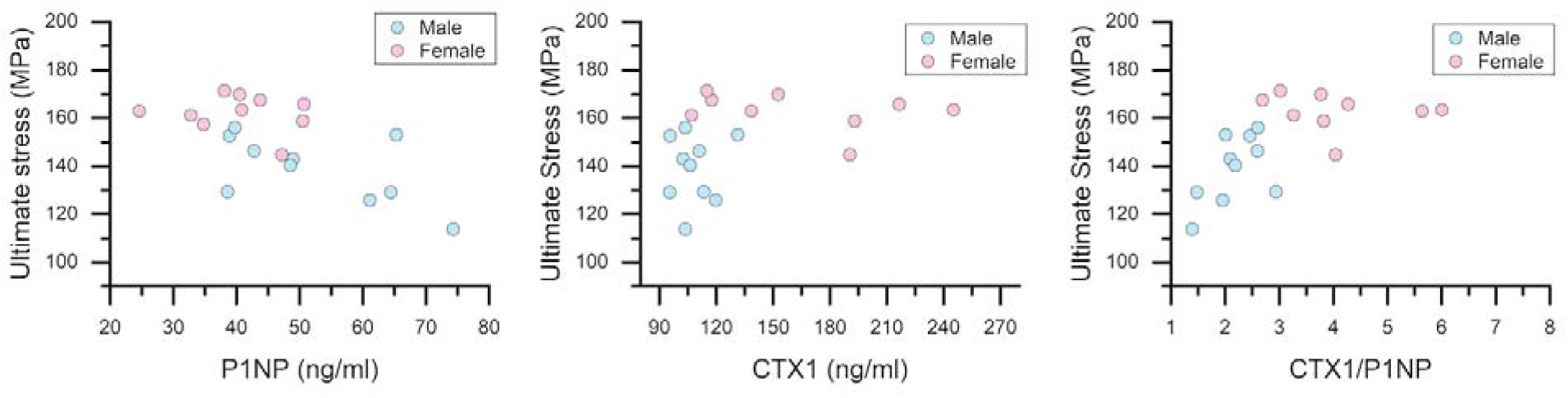
Correlation analysis between ultimate stress and serum biomarker concentrations. (A) For pooled male and female data, there is a significant negative correlation between P1NP and ultimate stress, and a significant positive correlation between ultimate stress and (B) CTX1 and (C) CTX1/P1NP.

## Supplementary Methods

**1. Description of Statistical Analyses performed for untargeted metabolomic profiling.**

Untargeted metabolomic profiling is performed to analyze all metabolite features detected in samples. This often yields large datasets, therefore, univariate, supervised, and unsupervised multivariate analyses can be performed to best visualize and narrow the dataset. A commonly used platform to analyze metabolomic data is MetaboAnalyst. Here, raw data can be normalized, transformed, and scaled. Specifically, normalization is recommended to adjust for systematic differences among samples. Data transformation, such as log or square root transformation, can be executed. Finally, scaling allows for adjustment of features based on dispersion of the variable of interest.

Depending on the number of experimental groups and comparisons of interest, various analyses, unsupervised and supervised, can be performed. Commonly, hierarchical cluster analysis (HCA), principal component analysis (PCA), partial least squares-discriminant analysis (PLS-DA), variable importance in projection (VIP) score, dendrograms, volcano plot analysis, t-test, and fold change can be utilized to visualize and analyze data.

Unsupervised multivariate statistical analyses that are commonly utilized to visualize metabolomic data are HCA and PCA. HCA builds tree structures based on data similarities. With this information, one can visualize metabolomic profiles, identify potential subgroups within experimental groups, and overall, visualize potential differences that may exist between groups of interest. To further examine metabolic data in an unsupervised way, PCA can be employed. In this statistical analysis, data is linearly transformed, and the large dataset is reduced into latent variables – principal components (PCs). PCs are a combination of metabolite features that explain the variability in the dataset. For example, a PC percentage of 18.2% suggests that 18.2% of the variability in the dataset is accounted for. Therefore, PCs allow researchers to gauge variability and distinguish the presence or absence of separation between groups.

A common next step following PCA is to perform PLS-DA, a supervised analysis. PLS-DA is like PCA as it helps visualize differences between cohorts but to do so, it uses a linear regression model and projects the predicted and observed variables. An extension of PLS-DA is VIP scores, which are useful to metabolomics because metabolites are scored based on their contribution to discrimination between groups. By performing these few statical methods, HCA, PCA, PLS-DA, and VIP scores initially provide an encompassing view of the similarities and differences between comparison groups of interest.

To further visualize data and begin to identify metabolite features, or groups of metabolite features, that are unique to one group, analyses like fold change and volcano plots can be performed. Implementing both tests when analyzing metabolomics data allows for metabolites that differ in intensity between groups to be elucidated.

Volcano plot is a common test applied to analyze metabolomic data because it shows statistical significance vs. magnitude of change. This visual displaying p-value and fold change together enables metabolite features that have large fold changes, are statistically significant, and belong to one experimental group but not the other to be identified. By identifying features that meet this criteria, biological relevance can be investigated. Specifically, differentially expressed metabolites can be subjected to pathway analysis using MetaboAnalyst’s MS Peaks to Pathways feature using the *mummichog* algorithm. This tool is able to take the metabolite compounds identified via the statistical tests describes to be associated with biological pathways. With this information, networks of functional cellular activity and differences in activity between groups can be investigated further.

## References

1. Berman AG, Damrath JG, Hatch J, Pulliam AN, Powell KM, Hinton M, et al. Effects of Raloxifene and tibial loading on bone mass and mechanics in male and female mice. Connect Tissue Res. Jan 2022;63(1):3–15. Epub 2021/01/12.

2. Glatt V, Canalis E, Stadmeyer L, Bouxsein ML. Age-related changes in trabecular architecture differ in female and male C57BL/6J mice. J Bone Miner Res. Aug 2007;22(8):1197–207. Epub 2007/05/10.

3. Liu Z, Solesio ME, Schaffler MB, Frikha-Benayed D, Rosen CJ, Werner H, et al. Mitochondrial Function Is Compromised in Cortical Bone Osteocytes of Long-Lived Growth Hormone Receptor Null Mice. J Bone Miner Res. Jan 2019;34(1):106–22. Epub 2018/09/15.

4. Meakin LB, Galea GL, Sugiyama T, Lanyon LE, Price JS. Age-related impairment of bones’ adaptive response to loading in mice is associated with sex-related deficiencies in osteoblasts but no change in osteocytes. J Bone Miner Res. Aug 2014;29(8):1859–71. Epub 2014/03/20.

5. Mumtaz H, Dallas M, Begonia M, Lara-Castillo N, Scott JM, Johnson ML, et al. Age-related and sex-specific effects on architectural properties and biomechanical response of the C57BL/6N mouse femur, tibia and ulna. Bone Rep. Jun 2020;12:100266. Epub 2020/05/19.

6. Oestreich AK, Onuzuriuke A, Yao X, Talton O, Wang Y, Pfeiffer FM, et al. Leprdb/+ Dams Protect Wild-type Male Offspring Bone Strength from the Detrimental Effects of a High-Fat Diet. Endocrinology. Aug 1 2020;161(8). Epub 2020/06/03.

7. Somerville JM, Aspden RM, Armour KE, Armour KJ, Reid DM. Growth of C57BL/6 mice and the material and mechanical properties of cortical bone from the tibia. Calcif Tissue Int. May 2004;74(5):469–75. Epub 2004/02/13.

8. Thiagarajan G, Begonia MT, Dallas M, Lara-Castillo N, Scott JM, Johnson ML. Determination of Elastic Modulus in Mouse Bones Using a Nondestructive Micro-Indentation Technique Using Reference Point Indentation. J Biomech Eng. Jul 1 2018;140(7). Epub 2018/05/26.

9. Vahidi G, Flook H, Sherk V, Mergy M, Lefcort F, Heveran CM. Bone biomechanical properties and tissue-scale bone quality in a genetic mouse model of familial dysautonomia. Osteoporos Int. Nov 2021;32(11):2335–46. Epub 2021/05/27.

10. Wallace JM, Rajachar RM, Allen MR, Bloomfield SA, Robey PG, Young MF, et al. Exercise-induced changes in the cortical bone of growing mice are bone- and gender-specific. Bone. Apr 2007;40(4):1120–7. Epub 2007/01/24.

11. Iwaniec UT, Wronski TJ, Liu J, Rivera MF, Arzaga RR, Hansen G, et al. PTH stimulates bone formation in mice deficient in Lrp5. J Bone Miner Res. Mar 2007;22(3):394–402. Epub 2006/12/07.

12. Johnson RW, McGregor NE, Brennan HJ, Crimeen-Irwin B, Poulton IJ, Martin TJ, et al. Glycoprotein130 (Gp130)/interleukin-6 (IL-6) signalling in osteoclasts promotes bone formation in periosteal and trabecular bone. Bone. Dec 2015;81:343–51. Epub 2015/08/11.

13. Martin SA, Philbrick KA, Wong CP, Olson DA, Branscum AJ, Jump DB, et al. Thermoneutral housing attenuates premature cancellous bone loss in male C57BL/6J mice. Endocr Connect. Nov 2019;8(11):1455–67. Epub 2019/10/08.

14. Mohammad KS, Chen CG, Balooch G, Stebbins E, McKenna CR, Davis H, et al. Pharmacologic inhibition of the TGF-beta type I receptor kinase has anabolic and anti-catabolic effects on bone. PLoS One. 2009;4(4):e5275. Epub 2009/04/10.

15. Pantschenko AG, Zhang W, Nahounou M, McCarthy MB, Stover ML, Lichtler AC, et al. Effect of osteoblast-targeted expression of bcl-2 in bone: differential response in male and female mice. J Bone Miner Res. Aug 2005;20(8):1414–29. Epub 2005/07/12.

16. Zanotti S, Canalis E. Notch1 and Notch2 expression in osteoblast precursors regulates femoral microarchitecture. Bone. May 2014;62:22–8. Epub 2014/02/11.

17. Creecy A, Uppuganti S, Girard MR, Schlunk SG, Amah C, Granke M, et al. The age-related decrease in material properties of BALB/c mouse long bones involves alterations to the extracellular matrix. Bone. Jan 2020;130:115126. Epub 2019/11/05.

18. Sherk VD, Heveran CM, Foright RM, Johnson GC, Presby DM, Ferguson VL, et al. Sex differences in the effect of diet, obesity, and exercise on bone quality and fracture toughness. Bone. Apr 2021;145:115840. Epub 2021/01/09.

19. Schlecht SH, Bigelow EM, Jepsen KJ. How Does Bone Strength Compare Across Sex, Site, and Ethnicity? Clin Orthop Relat Res. Aug 2015;473(8):2540–7. Epub 2015/03/06.

20. Sherk VD, Bemben DA. Age and sex differences in estimated tibia strength: influence of measurement site. J Clin Densitom. Apr-Jun 2013;16(2):196–203. Epub 2012/06/09.

21. Burr DB. Bone quality: understanding what matters. J Musculoskelet Neuronal Interact. Jun 2004;4(2):184–6. Epub 2004/12/24.

22. Seeman E. Bone quality: the material and structural basis of bone strength. J Bone Miner Metab. 2008;26(1):1–8. Epub 2007/12/21.

23. Dirckx N, Moorer MC, Clemens TL, Riddle RC. The role of osteoblasts in energy homeostasis. Nat Rev Endocrinol. Nov 2019;15(11):651–65. Epub 2019/08/30.

24. Yang J, Ueharu H, Mishina Y. Energy metabolism: A newly emerging target of BMP signaling in bone homeostasis. Bone. Sep 2020;138:115467. Epub 2020/06/09.

25. Guntur AR, Le PT, Farber CR, Rosen CJ. Bioenergetics during calvarial osteoblast differentiation reflect strain differences in bone mass. Endocrinology. May 2014;155(5):1589–95. Epub 2014/01/21.

26. Karthik V, Guntur AR. Energy Metabolism of Osteocytes. Curr Osteoporos Rep. Jun 12 2021. Epub 2021/06/13.

27. Tian L, Rosen CJ, Guntur AR. Mitochondrial Function and Metabolism of Cultured Skeletal Cells. Methods Mol Biol. 2021;2230:437–47. Epub 2020/11/17.

28. Wang K, Le L, Chun BM, Tiede-Lewis LM, Shiflett LA, Prideaux M, et al. A Novel Osteogenic Cell Line That Differentiates Into GFP-Tagged Osteocytes and Forms Mineral With a Bone-Like Lacunocanalicular Structure. J Bone Miner Res. Jun 2019;34(6):979–95. Epub 2019/03/19.

29. Li B, Lee WC, Song C, Ye L, Abel ED, Long F. Both aerobic glycolysis and mitochondrial respiration are required for osteoclast differentiation. FASEB J. Aug 2020;34(8):11058–67. Epub 2020/07/07.

30. Li X, Wang X, Zhang C, Wang J, Wang S, Hu L. Dysfunction of metabolic activity of bone marrow mesenchymal stem cells in aged mice. Cell Prolif. Jan 27 2022:e13191. Epub 2022/01/29.

31. Shum LC, White NS, Nadtochiy SM, Bentley KL, Brookes PS, Jonason JH, et al. Cyclophilin D Knock-Out Mice Show Enhanced Resistance to Osteoporosis and to Metabolic Changes Observed in Aging Bone. PLoS One. 2016; 11(5):e0155709. Epub 2016/05/18.

32. Zhao H, Li X, Zhang D, Chen H, Chao Y, Wu K, et al. Integrative Bone Metabolomics-Lipidomics Strategy for Pathological Mechanism of Postmenopausal Osteoporosis Mouse Model. Sci Rep. Nov 7 2018;8(1):16456. Epub 2018/11/09.

33. Travlos GS. Normal structure, function, and histology of the bone marrow. Toxicol Pathol. 2006;34(5):548–65. Epub 2006/10/28.

34. Bonewald LF. The amazing osteocyte. J Bone Miner Res. Feb 2011;26(2):229–38. Epub 2011/01/22.

35. Schaffler MB, Cheung WY, Majeska R, Kennedy O. Osteocytes: master orchestrators of bone. Calcif Tissue Int. Jan 2014;94(1):5–24. Epub 2013/09/18.

36. Bouxsein ML, Boyd SK, Christiansen BA, Guldberg RE, Jepsen KJ, Muller R. Guidelines for assessment of bone microstructure in rodents using microcomputed tomography. J Bone Miner Res. Jul 2010;25(7):1468–86. Epub 2010/06/10.

37. Turner CH, Burr DB. Basic biomechanical measurements of bone: a tutorial. Bone. Jul-Aug 1993;14(4):595–608. Epub 1993/07/01.

38. Ritchie RO, Koester KJ, Ionova S, Yao W, Lane NE, Ager JW, 3rd. Measurement of the toughness of bone: a tutorial with special reference to small animal studies. Bone. Nov 2008;43(5):798–812. Epub 2008/07/24.

39. Hahn AK, Wallace CW, Welhaven HD, Brooks E, McAlpine M, Christiansen BA, et al. The microbiome mediates epiphyseal bone loss and metabolomic changes after acute joint trauma in mice. Osteoarthritis Cartilage. Jun 2021;29(6):882–93. Epub 2021/03/22.

40. Carlson AK, Rawle RA, Wallace CW, Adams E, Greenwood MC, Bothner B, et al. Global metabolomic profiling of human synovial fluid for rheumatoid arthritis biomarkers. Clin Exp Rheumatol. May-Jun 2019;37(3):393–9. Epub 2019/01/09.

41. Bar N, Korem T, Weissbrod O, Zeevi D, Rothschild D, Leviatan S, et al. A reference map of potential determinants for the human serum metabolome. Nature. Dec 2020;588(7836):135–40. Epub 2020/11/13.

42. Carlson AK, Rawle RA, Adams E, Greenwood MC, Bothner B, June RK. Application of global metabolomic profiling of synovial fluid for osteoarthritis biomarkers. Biochem Biophys Res Commun. May 5 2018;499(2):182–8. Epub 2018/03/20.

43. Carlson AK, Rawle RA, Wallace CW, Brooks EG, Adams E, Greenwood MC, et al. Characterization of synovial fluid metabolomic phenotypes of cartilage morphological changes associated with osteoarthritis. Osteoarthritis Cartilage. Aug 2019;27(8):1174–84. Epub 2019/04/28.

44. Hugle T, Kovacs H, Heijnen IA, Daikeler T, Baisch U, Hicks JM, et al. Synovial fluid metabolomics in different forms of arthritis assessed by nuclear magnetic resonance spectroscopy. Clin Exp Rheumatol. Mar-Apr 2012;30(2):240–5. Epub 2012/03/14.

45. Jutila AA, Zignego DL, Hwang BK, Hilmer JK, Hamerly T, Minor CA, et al. Candidate mediators of chondrocyte mechanotransduction via targeted and untargeted metabolomic measurements. Arch Biochem Biophys. Mar 1 2014;545:116–23. Epub 2014/01/21.

46. Kim S, Hwang J, Kim J, Ahn JK, Cha HS, Kim KH. Metabolite profiles of synovial fluid change with the radiographic severity of knee osteoarthritis. Joint Bone Spine. Oct 2017;84(5):605–10. Epub 2016/07/28.

47. Kim S, Hwang J, Xuan J, Jung YH, Cha HS, Kim KH. Global metabolite profiling of synovial fluid for the specific diagnosis of rheumatoid arthritis from other inflammatory arthritis. PLoS One. 2014;9(6):e97501. Epub 2014/06/03.

48. Lamers RJ, van Nesselrooij JH, Kraus VB, Jordan JM, Renner JB, Dragomir AD, et al. Identification of an urinary metabolite profile associated with osteoarthritis. Osteoarthritis Cartilage. Sep 2005;13(9):762–8. Epub 2005/06/14.

49. Loeser RF, Pathmasiri W, Sumner SJ, McRitchie S, Beavers D, Saxena P, et al. Association of urinary metabolites with radiographic progression of knee osteoarthritis in overweight and obese adults: an exploratory study. Osteoarthritis Cartilage. Aug 2016;24(8):1479–86. Epub 2016/03/26.

50. McCutchen CN, Zignego DL, June RK. Metabolic responses induced by compression of chondrocytes in variable-stiffness microenvironments. J Biomech. Nov 7 2017;64:49–58. Epub 2017/10/08.

51. Salinas D, Mumey BM, June RK. Physiological dynamic compression regulates central energy metabolism in primary human chondrocytes. Biomech Model Mechanobiol. Feb 2019;18(1):69–77. Epub 2018/08/12.

52. Yang G, Zhang H, Chen T, Zhu W, Ding S, Xu K, et al. Metabolic analysis of osteoarthritis subchondral bone based on UPLC/Q-TOF-MS. Anal Bioanal Chem. Jun 2016;408(16):4275–86. Epub 2016/04/15.

53. Zhang W, Likhodii S, Zhang Y, Aref-Eshghi E, Harper PE, Randell E, et al. Classification of osteoarthritis phenotypes by metabolomics analysis. BMJ Open. Nov 19 2014;4(11):e006286. Epub 2014/11/21.

54. Zhang W, Sun G, Likhodii S, Liu M, Aref-Eshghi E, Harper PE, et al. Metabolomic analysis of human plasma reveals that arginine is depleted in knee osteoarthritis patients. Osteoarthritis Cartilage. May 2016;24(5):827–34. Epub 2015/12/29.

55. Zignego DL, Hilmer JK, June RK. Mechanotransduction in primary human osteoarthritic chondrocytes is mediated by metabolism of energy, lipids, and amino acids. J Biomech. Dec 16 2015;48(16):4253–61. Epub 2015/11/18.

56. Guma M, Tiziani S, Firestein GS. Metabolomics in rheumatic diseases: desperately seeking biomarkers. Nat Rev Rheumatol. May 2016;12(5):269–81. Epub 2016/03/05.

57. Kanehisa M, Sato Y, Kawashima M. KEGG mapping tools for uncovering hidden features in biological data. Protein Sci. Jan 2022;31(1):47–53. Epub 2021/08/24.

58. Hawkes CP, Mostoufi-Moab S. Fat-bone interaction within the bone marrow milieu: Impact on hematopoiesis and systemic energy metabolism. Bone. Feb 2019;119:57–64. Epub 2018/03/20.

59. Li M, Zhai L, Wei W, Dong J. Effect of Methionine Restriction on Bone Density and NK Cell Activity. Biomed Res Int. 2016;2016:3571810. Epub 2016/11/25.

60. Parkhitko AA, Jouandin P, Mohr SE, Perrimon N. Methionine metabolism and methyltransferases in the regulation of aging and lifespan extension across species. Aging Cell. Dec 2019;18(6):e13034. Epub 2019/08/29.

61. Baines M, Kredan MB, Davison A, Higgins G, West C, Fraser WD, et al. The association between cysteine, bone turnover, and low bone mass. Calcif Tissue Int. Dec 2007;81(6):450–4. Epub 2007/12/07.

62. Ostrakhovitch EA, Tabibzadeh S. Homocysteine and age-associated disorders. Ageing Res Rev. Jan 2019;49:144–64. Epub 2018/11/06.

63. Tetlow LC, Woolley DE. Histamine, histamine receptors (H1 and H2), and histidine decarboxylase expression by chondrocytes of osteoarthritic cartilage: an immunohistochemical study. Rheumatol Int. Dec 2005;26(2):173–8. Epub 2005/06/30.

64. Tetlow LC, Woolley DE. Histamine stimulates the proliferation of human articular chondrocytes in vitro and is expressed by chondrocytes in osteoarthritic cartilage. Ann Rheum Dis. Oct 2003;62(10):991–4. Epub 2003/09/16.

65. Xia MF, Lin HD, Yan HM, Bian H, Chang XX, Zhang LS, et al. The association of liver fat content and serum alanine aminotransferase with bone mineral density in middle-aged and elderly Chinese men and postmenopausal women. J Transl Med. Jan 13 2016;14:11. Epub 2016/01/23.

66. Bellissimo MP, Roberts JL, Jones DP, Liu KH, Taibl KR, Uppal K, et al. Metabolomic Associations with Serum Bone Turnover Markers. Nutrients. Oct 16 2020;12(10). Epub 2020/10/22.

67. Consoli A, Nurjhan N, Reilly JJ, Jr., Bier DM, Gerich JE. Contribution of liver and skeletal muscle to alanine and lactate metabolism in humans. Am J Physiol. Nov 1990;259(5 Pt 1):E677–84. Epub 1990/11/11.

68. Skerry TM. The role of glutamate in the regulation of bone mass and architecture. J Musculoskelet Neuronal Interact. Apr-Jun 2008;8(2):166–73. Epub 2008/07/16.

69. Mason DJ, Suva LJ, Genever PG, Patton AJ, Steuckle S, Hillam RA, et al. Mechanically regulated expression of a neural glutamate transporter in bone: a role for excitatory amino acids as osteotropic agents? Bone. Mar 1997;20(3):199–205. Epub 1997/03/01.

70. Forrest CM, Mackay GM, Oxford L, Stoy N, Stone TW, Darlington LG. Kynurenine pathway metabolism in patients with osteoporosis after 2 years of drug treatment. Clin Exp Pharmacol Physiol. Nov 2006;33(11):1078–87. Epub 2006/10/18.

71. Fatokun AA, Stone TW, Smith RA. Hydrogen peroxide-induced oxidative stress in MC3T3-E1 cells: The effects of glutamate and protection by purines. Bone. Sep 2006;39(3):542–51. Epub 2006/04/18.

72. Chenu C, Serre CM, Raynal C, Burt-Pichat B, Delmas PD. Glutamate receptors are expressed by bone cells and are involved in bone resorption. Bone. Apr 1998;22(4):295–9. Epub 1998/04/29.

73. Kushwaha P, Wolfgang MJ, Riddle RC. Fatty acid metabolism by the osteoblast. Bone. Oct 2018;115:8–14. Epub 2017/09/03.

74. Niemeier A, Niedzielska D, Secer R, Schilling A, Merkel M, Enrich C, et al. Uptake of postprandial lipoproteins into bone in vivo: impact on osteoblast function. Bone. Aug 2008;43(2):230–7. Epub 2008/06/10.

75. Catherwood BD, Addison J, Chapman G, Contreras S, Lorang M. Growth of rat osteoblast-like cells in a lipid-enriched culture medium and regulation of function by parathyroid hormone and 1,25-dihydroxyvitamin D. J Bone Miner Res. Aug 1988;3(4):431–8. Epub 1988/08/01.

76. Kim SP, Li Z, Zoch ML, Frey JL, Bowman CE, Kushwaha P, et al. Fatty acid oxidation by the osteoblast is required for normal bone acquisition in a sex- and diet-dependent manner. JCI Insight. Aug 17 2017;2(16). Epub 2017/08/18.

77. Bartelt A, Koehne T, Todter K, Reimer R, Muller B, Behler-Janbeck F, et al. Quantification of Bone Fatty Acid Metabolism and Its Regulation by Adipocyte Lipoprotein Lipase. Int J Mol Sci. Jun 13 2017;18(6). Epub 2017/06/14.

78. van Gastel N, Stegen S, Eelen G, Schoors S, Carlier A, Daniels VW, et al. Lipid availability determines fate of skeletal progenitor cells via SOX9. Nature. Mar 2020;579(7797):111–7. Epub 2020/02/28.

79. Ishii KA, Fumoto T, Iwai K, Takeshita S, Ito M, Shimohata N, et al. Coordination of PGC-1beta and iron uptake in mitochondrial biogenesis and osteoclast activation. Nat Med. Mar 2009;15(3):259–66. Epub 2009/03/03.

80. Wei W, Wang X, Yang M, Smith LC, Dechow PC, Sonoda J, et al. PGC1beta mediates PPARgamma activation of osteoclastogenesis and rosiglitazone-induced bone loss. Cell Metab. Jun 9 2010;11(6):503–16. Epub 2010/06/04.

81. Zhou J, Ye S, Fujiwara T, Manolagas SC, Zhao H. Steap4 plays a critical role in osteoclastogenesis in vitro by regulating cellular iron/reactive oxygen species (ROS) levels and cAMP response element-binding protein (CREB) activation. J Biol Chem. Oct 18 2013;288(42):30064–74. Epub 2013/08/31.

82. Hernandez CJ. How can bone turnover modify bone strength independent of bone mass? Bone. Jun 2008;42(6):1014–20. Epub 2008/04/01.

83. Le Floc’h N, Otten W, Merlot E. Tryptophan metabolism, from nutrition to potential therapeutic applications. Amino Acids. Nov 2011;41(5):1195–205. Epub 2010/09/28.

84. Michalowska M, Znorko B, Kaminski T, Oksztulska-Kolanek E, Pawlak D. New insights into tryptophan and its metabolites in the regulation of bone metabolism. J Physiol Pharmacol. Dec 2015;66(6):779–91. Epub 2016/01/16.

85. Sainio EL, Pulkki K, Young SN. L-Tryptophan: Biochemical, nutritional and pharmacological aspects. Amino Acids. Mar 1996;10(1):21–47. Epub 1996/03/01.

86. Forrest CM, Kennedy A, Stone TW, Stoy N, Darlington LG. Kynurenine and neopterin levels in patients with rheumatoid arthritis and osteoporosis during drug treatment. Adv Exp Med Biol. 2003;527:287–95. Epub 2004/06/23.

87. El Refaey M, Watkins CP, Kennedy EJ, Chang A, Zhong Q, Ding KH, et al. Oxidation of the aromatic amino acids tryptophan and tyrosine disrupts their anabolic effects on bone marrow mesenchymal stem cells. Mol Cell Endocrinol. Jul 15 2015;410:87–96. Epub 2015/02/01.

88. Siddiqui A, Ceppi P. A non-proliferative role of pyrimidine metabolism in cancer. Mol Metab. May 2020;35:100962. Epub 2020/04/04.

